# Decoding the Epitranscriptional Landscape from Native RNA Sequences

**DOI:** 10.1101/487819

**Authors:** Thidathip Wongsurawat, Piroon Jenjaroenpun, Trudy M. Wassenaar, Taylor D Wadley, Visanu Wanchai, Nisreen S. Akel, Aime T. Franco, Michael L. Jennings, David W. Ussery, Intawat Nookaew

**Affiliations:** Department of Biomedical Informatics, College of Medicine, University of Arkansas for Medical Sciences, Little Rock, AR 72205, USA; Molecular Microbiology and Genomics Consultants, Zotzenheim, Germany; Department of Physiology and Biophysics, College of Medicine, The University of Arkansas for Medical Sciences, Little Rock, AR 72205, USA

**Author notes:** **Corresponding author**: Intawat Nookaew. Department of Biomedical Informatics, College of Medicine, University of Arkansas for Medical Sciences, 4301 West Markham Street, Slot 782, Little Rock, AR 72205, USA. These authors contributed equally to this work (Thidathip Wongsurawat, Piroon Jenjaroenpun).

## Abstract

Sequencing of native RNA and corresponding cDNA was performed using Oxford Nanopore Technology. The % Error of Specific Bases (%ESB) was higher for native RNA than for cDNA, which enabled detection of ribonucleotide modification sites. Based on %ESB differences of the two templates, a bioinformatic tool ELIGOS was developed and applied to rRNAs of *E. coli,* yeast and human cells. ELIGOS captured 91%, 95%, ∼75%, respectively, of the known variety of RNA methylation sites in these rRNAs. Yeast transcriptomes from different growth conditions were also compared, which identified an association between metabolic adaptation and inferred RNA modifications. ELIGOS was further applied to human transcriptome datasets, which identified the well-known DRACH motif containing N6-methyadenine being located close to 3’-untranslated regions of mRNA. Moreover, the RNA G-quadruplex motif was uncovered by ELIGOS. In summary, we have developed an experimental method coupled with bioinformatic software to uncover native RNA modifications and secondary-structures within transcripts.

## MAIN TEXT

The transcriptome is the collection of all RNA molecules present in a given cell that can be determined by high-throughput techniques, such as microarray analysis or RNA sequencing (RNA-seq) methods ^1^. RNA-seq using next-generation sequencing (NGS) techniques has been replacing microarray analysis, since the former is able to detect novel or unknown transcripts. Further, NGS enables transcriptome analysis with a higher dynamic range of expression levels than for microarrays ^2^. With improved sample preparation methods and reduced sequencing costs, RNA-seq by NGS has become the method of choice to study transcriptomes.

The length of sequence reads generated with most NGS platforms range from 35 nt up to about 500 nt, so that single reads rarely cover a complete transcript. Accurate alignment and assembly of such short sequences depends on availability of a reference genome, and the identification of spliced isoforms or gene-fusion transcripts remains a challenge ^3^. Further, methods depending on reverse transcription (RT) of RNA and amplification may introduce biases and artifacts ^4^. These shortcomings can be overcome by directly sequencing native RNA molecules using technologies such as the Oxford Nanopore Technologies (ONT) platform. Direct RNA sequencing without amplification (dRNA-seq) is able to generate long reads, typically covering the full length of a transcript ^5^. The method can accurately quantify transcripts in order to analyze differential gene expression with a dynamic range comparable to traditional RNA-seq derived from short read sequencing, while it enables accurate identification of the structure and boundaries of transcripts including spliced products ^6^.

An additional advantage of dRNA-seq is the detection of transcriptional modifications inferred from the current signal as the RNA molecule passes a nanopore: modified RNA molecules cause a characteristic current blockade, enabling simultaneous detection of diverse modifications such as 5-methylcytosine (m5C) or 6-methyladenine (m6A) ^5, 7, 8^. Presently, over 170 different types of RNA modifications have been described within the prokaryote and eukaryote kingdoms, which are collected in various databases ^9, 10, 11^. High throughput sequencing coupled with methods to specifically enrich RNA modification products create the possibility to study the epigenetics of RNA and describe the ‘epitranscriptome’, a term introduced in 2012 ^12^. However, these methods are labor intensive and may introduce experimental artifacts or biased results, and they suffer from a relatively high false positive rate ^13^. Moreover, the transcriptome-wide approach nowadays can only identify only a dozen from 170 known different types of RNA modifications because limitation of available specific antibodies or chemical treatmnets^14^. Alternatively, using the traditional approach of LC–MS/MS can identify several types of modification however, the approach has limitations to identify the transcript that contains modifications and their position of modifications ^14^.

ONT sequencing also has certain disadvantages, the main one being a relatively high error rate. Translation of the obtained electrical current signals into specific bases relies on either trained hidden Markov or neural network models ^15^. The accuracy of individual DNA reads is currently around 90% on average ^15^; and we typically experience an accuracy of about 88% in RNA reads ^6^. The most commonly encountered errors are related to presence of homopolymers, base modifications, nucleic acid damage and structural features of the nucleic acid molecules.

It is known that Reverse Transcriptase can ignore modifications of the RNA template to produce cDNA devoid of modification information ^16^. We anticipated that the ONT sequencing signals obtained from cDNA and those derived from the same RNA molecules by dRNA-seq could be used to filter out systematic noise from data to detect locations of possible RNA modifications. To test this, we used *in vitro* transcripts of a luciferase gene produced with and without incorporation of 5-methoxy-uridine (5moU). By comparison of the resultant dRNA-seq data of unmodified and modified RNA with those obtained from direct cDNA sequencing (dcDNA-seq), we were able to filter out signals that were most likely due to presence of modified bases.

The software tool “Epitranscriptional Landscape Inferring from Glitches of ONT Signals” (ELIGOS) was developed to predict the presence of modified bases from a comparison of dRNA-seq and dcDNA-seq data, and the output of this tool was verified with ribosomal RNA sequences from yeast, bacteria (*Escherichia coli*) and human cells, after which the procedure was used to map the yeast transcriptome. Transcripts of *Saccharomyces cerevisiae* strain CEN.PK113-7D were compared for cells cultured in minimal medium in presence of glucose and under glucose depletion, and these were compared to transcripts of *S. cerevisiae* strain DBY746 grown in rich medium. The comparison was extended to the transcriptome of a human cell line, from which hyper-modified transcripts were identified. The implications of this novel approach to investigate the epitranscriptome of cells are discussed.

## Results

### Distinguishing modified RNA bases from sequencing errors

The nanopore sequencing signal of RNA can be affected by three-dimensional structures of the RNA template, as well as by presence of modified ribonucleotides, both of which can lead to sequencing errors. Since modified bases are absent when RNA is converted into cDNA, we anticipated that an in-depth analysis of sequencing errors for both types of templates might be able to differentiate between the presence of modified bases and stochastic errors. In a pilot experiment, we mimicked post-transcriptional modifications of RNA by *in vitro* incorporation of 5-methoxy-uridine (5moU) into transcripts of a luciferase gene. Sequencing signals were compared for this modified mRNA (dRNA^O^), the corresponding dcDNA (dcDNA^O^), and from dRNA sequences obtained with unmodified uridine (dRNA^U^).

Figure 1 shows that *in vitro* incorporation of 5moU resulted in dRNA^O^ reads with significantly higher % Error at Specific Bases (%ESB, defined as described in the methods) than dcDNA^O^ (p-value 8.3e^-94^) or dRNA^U^ (p-value 4.6e^-118^). Notably, for values up to approximately 25%, the distributions of %ESB for both dRNA^U^ and dcDNA^O^ were overlapping and higher than those for dRNA^O^, but for values above 25%, dRNA^O^ reported significantly higher %ESB (Figure 1A). We interpret this to mean that below 25% ESB the error rate was mostly due to random noise, but the increased %ESB of dRNA above that cut-off might reflect a biological signal, possibly (but not exclusively) related to presence of modified bases that can be used to distinguish true signal from background noise.

**Figure 1.**
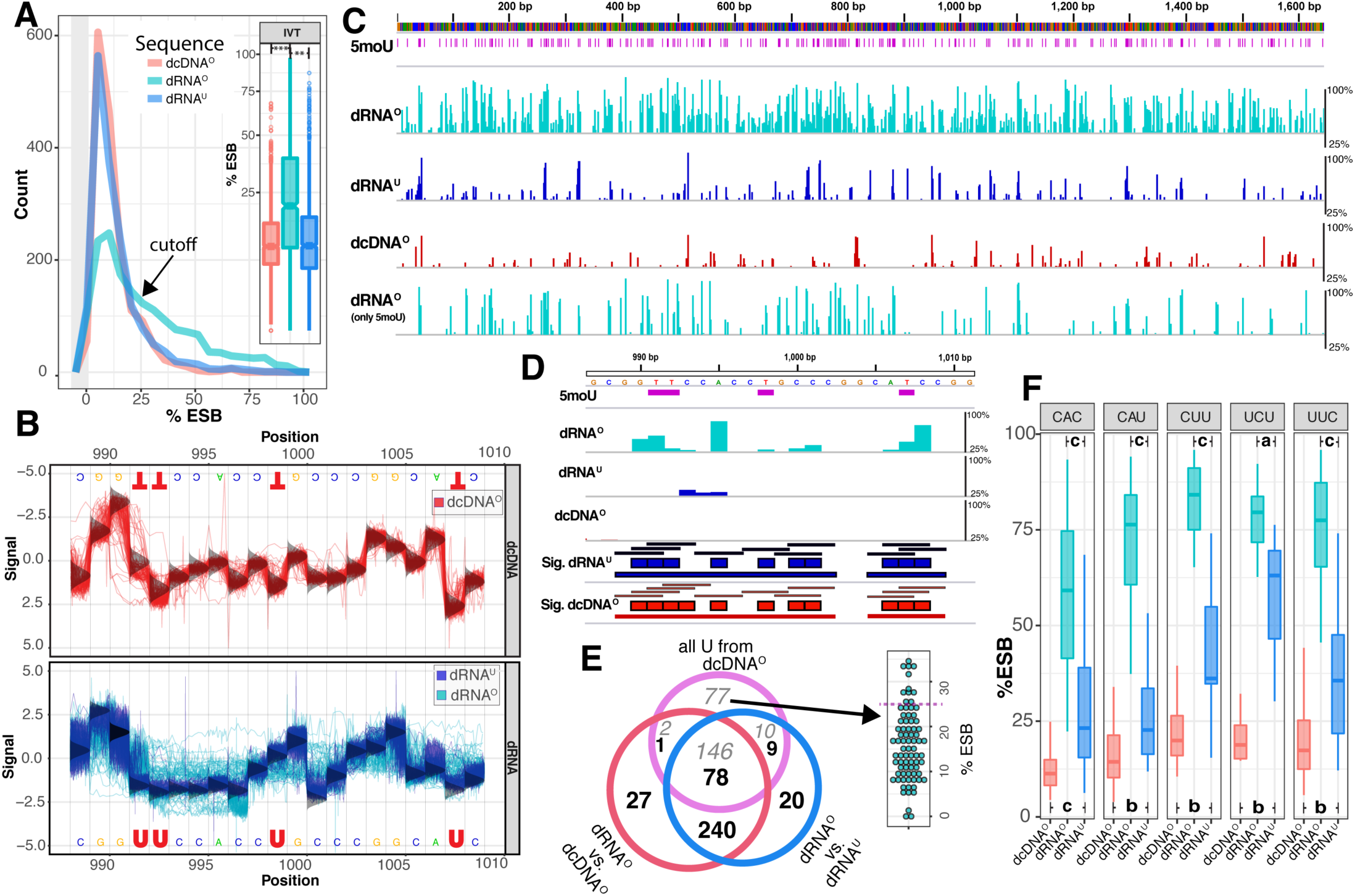
Direct sequencing of *in vitro* transcripts of the luciferase gene with and without incorporation of 5-methoxy-uridine. **(A)** The distribution of the percentage <u>E</u>rror at a <u>S</u>pecific <u>B</u>ase (%ESB) for dRNA^O^ differs significantly from that of dcDNA^O^ and dRNA^U^, with ** P<e^-60^, ***<e^-100^. The black arrow indicates at which frequency of %ESB higher values are found in dRNA^O^ than in the other two templates. The thick gray area to the left of the plot represents the histogram of the first bin around zero. **(B)** Re-squiggled signal plots of a selected region obtained with dcDNA^O^ template (top), and overlaid signals obtained with dRNA^U^ (blue) and dRNA^O^ (cyan) (bottom). The vertical, bell-shaped curves at each base position represent the distribution of the standard canonical model signals for either template. **C)** Position-specific %ESB passing the 25% cutoff for (from top downwards) dRNA^O^, dRNA^U^ and dcDNA^O^. The bottom line presents %ESB of dRNA^O^ only for positions where U is present. The positions of all uridines are shown in magenta below the colored sequence line. **(D**) Locus determination based on differential %ESB positions and merging of adjacent signals. From the top: 5moU positions shown as magenta bars; %ESB of dRNA^O^ sequences shown as cyan bars; %ESB of dRNA^O^ sequences shown as blue bars; dcDNA^O^ lane indicating absence of % ESB that pass the cutoff of 25 %; Sig. dRNA^U^ and Sig. dcDNA^O^ lanes illustrating the differential %ESB detected when comparing dRNA^O^ with dRNA^U^ (blue) or dcDNA^O^ (red), respectively. The middle colored blocks represent the differential ESB positions, the thinner black bars above them represent the locus extension with flanking bases on both sides, while the thin bars below the colored blocks represent the resultant merged loci. **(E)** Venn diagram of loci (black numbers) identified by differential %ESB values of dRNA^O^ compared to dcDNA^O^ (red circle), or compared to dRNA^U^ (blue circle). The numbers of all uridine positions are given in gray. To the right of the Venn diagram is the %ESB distribution shown for the 77 uridine positions not overlapping with the other two datasets. **(F)** Artifactual differential %ESB signals are sequence-dependent. The %ESB values of five identified triplets that differed significantly between dRNA^U^ and dcDNA^O^ or dRNA^O^ (*a*: p<0.05, *b*: p<e^-3^ and *c*: p<e^-8^ as derived from Wilcoxson’s rank sum test).

To illustrate the effect on recorded signals when modified bases are present, in Figure 1B the re-squiggled signals are compared for a small region (position 989-1009) of the luciferase gene containing four uracil bases in three loci. The sequence signals obtained with dcDNA^O^ (Figure 1B, in red) or from directly sequencing RNA^U^ (in blue) matched those of the theoretical canonical signal model for DNA. In contrast, the re-squiggle signals of dRNA^O^ containing modified uridine were altered compared to the canonical RNA signals (Figure 1B, in cyan). Thus, presence of 5moU bases most likely caused some of the observed perturbations, while an RT step removed this effect. Not only the 5moU sites, but also bases in their vicinity produced dramatically perturbed signals in dRNA^O^, for instance at position 997 (Figure 1B). This has a direct impact on the accuracy of base calling. Note, that base calling is typically performed on a window of 5-mers, so that any effect due to presence of a modified base can affect the signal of bases in its direct vicinity.

The positions for which %ESB exceeded the cutoff of 25% were recorded for the complete dRNA^O^ template, as well as for the templates dRNA^U^ and dcDNA^O^ (Figure 1C). High %ESB values were more frequently obtained with dRNA^O^ template than with either dRNA^U^ or dcDNA^O^. Further, positions where 5moU was present frequently produced higher %ESB. We also recorded >25% ESB values for some positions where other bases were present, and not all positions with 5moU did increase the %ESB in the dRNA^O^ reads. Some of the observed errors are due to the reduced speed of nucleotide translocation through the nanopore, causing a ‘glitch’ in the corresponding output. In a number of cases, high %ESB coincided with presence of homopolymeric stretches (Supplementary Figure S1). Although such signals are not easily distinguishable from signals due to base modifications, homopolymeric stretches can be readily identified from the sequence. Further, elevated %ESB values observed in both dRNA^U^ and dcDNA^O^ are more likely to be caused by structural features irrespective to presence of modified bases, and these should ideally be removed from the data.

To this extent we developed a bioinformatics software tool, ELIGOS, that determines differential %ESB positions between dRNA and a reference sequence (either cDNA or non-modified RNA of the same sequence). We used a cut-off for an odds ratio of ≥2 and adjusted p-values <1e^-50^ to identify differential %ESB positions. The optimal %ESB cutoff was determined as 25% based on a loss-gain analysis using a 20-30% range, as shown in Figure S2.

Since the presence of a methylated base can influence the differential %ESB of adjacent positions, flanking bases should also be considered (as exemplified in Figure 1B where the signal of bases in the vicinity of 5moU was sometimes altered). Thus, we first recorded all positions for which the %ESB between the dRNA^O^ signal and the reference signals differed. These positions were then extended to the flanking bases positioned directly 5’ and 3’ to produce triplet loci. These triplets were individually assessed, unless two recorded triplets overlapped or were direct neighbors, in which case their locus was extended, as shown in the example of Figure 1D.

A total of 346 and 347 loci with differential %ESB were identified in the luciferase transcript using dcDNA^O^ and dRNA^U^ as the reference, respectively. These loci overlapped in 318 cases. Since for the *in vitro* transcripts the exact positions of all methylated bases were known (*i.e*., all uridine was 5moU), their positions were compared to the identified loci to assess how well these matched with presence of methylated bases (Figure 1E). We found that 78 identified loci contained at least one 5moU (in total these covered 146 5moU bases). The differential %ESB values that had identified these loci were likely caused by presence of the modified 5MoU bases, while potential loci not containing uracil may have been caused by features unrelated to base modification.

Ideally, direct sequencing of unmodified RNA as a reference for comparison would be best. However, this is not practical for most biological systems, where in most cases dcDNA and native RNA are available. If dcDNA were the only available reference, our findings would be similar, since only one locus identified with that reference did not match the findings obtained with dRNA^U^. We take this as evidence that the approach to compare differential %ESB values obtained from cDNA and modified RNA can indeed identify the presence of modified bases. We found 77 moU positions that did not produce elevated differentiated %ESB values in dRNA-seq signals when compared to dcDNA^O^ or dRNA^U^; these produced %ESB values <25% in 67 of 77 cases (87%). These findings illustrate the limitation in case of a heavily modified synthetic RNA template, that contained the maximum fraction of modified uridine bases. We then continued to our investigations with natural modified RNA that normally has much lesser fraction of RNA modifications.

### Presence of artifactual specific triplets in dRNA-seq data of IVT

We next checked whether any of the ELIGOS output data were caused by sequence-dependent artifacts, by comparing the %ESB of all possible 52 triplets present in the luciferase gene in the dcDNA^O^, dRNA^U^ and dRNA^O^ data (see Supplementary Table S1 for details). This identified five triplets producing not only significantly higher %ESB values over the 25% threshold for dRNA^O^ but also for unmodified dRNA^U^, when compared to dcDNA^O^. These were CAC, CAU, CUU, UCU and UUC (Figure 1F). The differential %ESB for CAC could not be caused by presence of 5moU, so this was more likely due to a structural feature caused by this combination of nucleotides. For all these five triplets an inherit signal amplification of dRNA-seq was present that needs to be corrected for when cDNA is solely used as the reference for differential % ESB position. The remaining 47 triplets did not result in strongly elevated %ESB signals for dRNA^U^ compared to dcDNA (see Supplementary Table S1), confirming that using dRNA^U^ as the reference for differential %ESB position determination is a valid approach, while after subtraction of systematic errors from signals truly due to base modification, dcDNA sequences can be used as a reference.

### Evaluation of ELIGOS for prediction of modified rRNA bases

The validity of ELIGOS predictions was tested for sequencing data obtained with ribosomal RNA (rRNA) from *S. cerevisiae, E. coli* and a human cell line, as the presence of modified bases and secondary structures in these RNA molecules has been extensively characterized. Total RNA was sequenced by dRNA-seq and dcDNA-seq, after which signals for the combined rRNA genes were extracted from the data. As observed with the *in vitro* transcripts, dRNA data for the rRNA produced significantly higher %ESB values than dcDNA, for all three organisms, with p-values of 2.5e^-118^, 4.9e^-40^, and 3.0e^-50^, for yeast, *E. coli* and human cells, respectively (Figure 2A).

**Figure 2.**
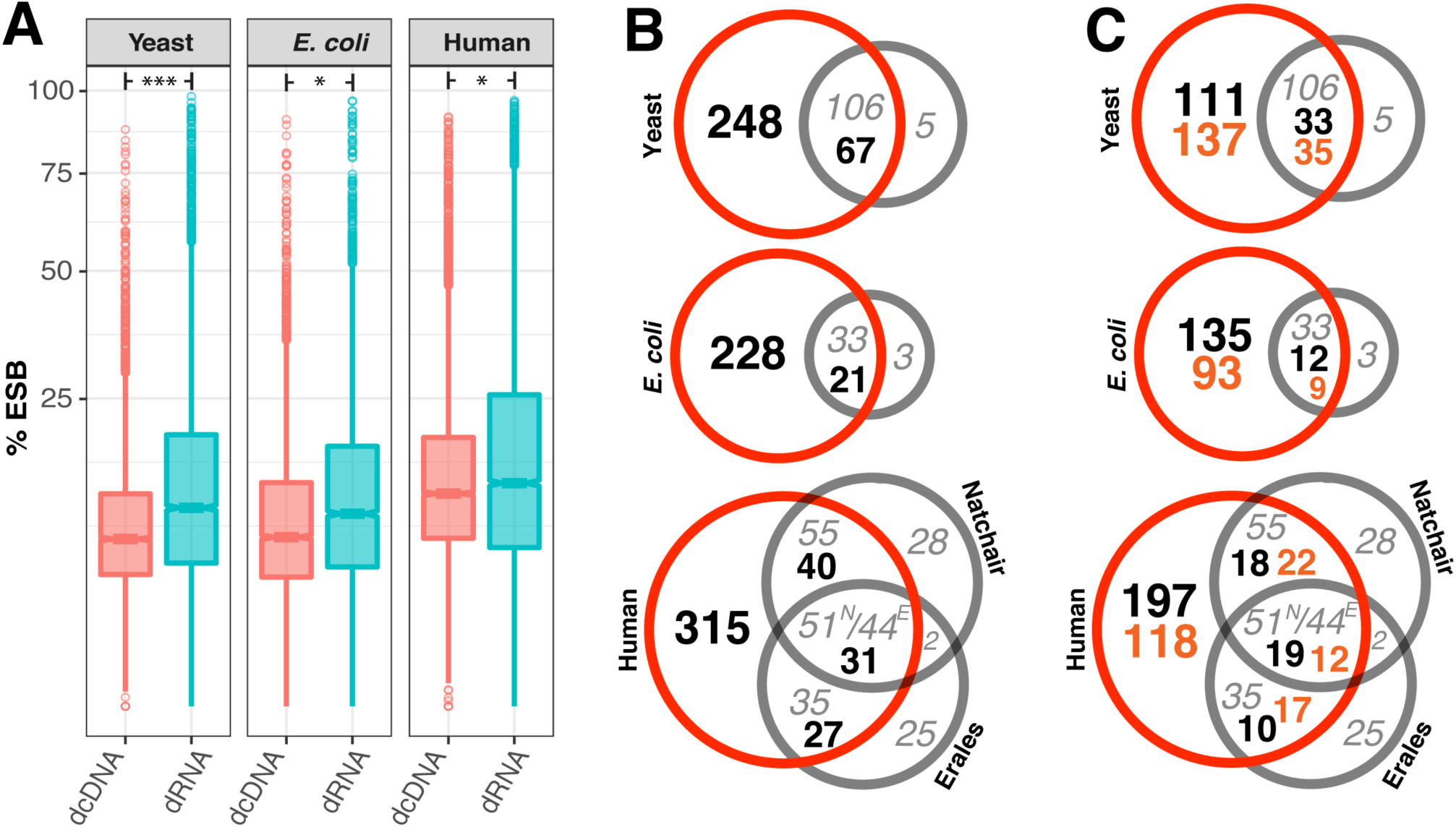
Direct sequencing of native rRNA and corresponding cDNA of yeast, *E. coli* and human cells. **(A)** The %ESB for dRNA differs significantly from that of dcDNA with * p<e^-30^, *** p<e-^100^ derived from Student’s *t*-test. **(B)** Venn diagrams showing in red circles ELIGOS-predicted loci (black numbers) and individual base positions (gray numbers) overlapping with described methylation sites (gray circles), for the three species. The human cell line data were compared to known methylation information retrieved from Natchair *et al*. ^21^ (superscript N in central interception) and Erales et al. ^20^ (superscript E). **(C)** The same Venn diagrams, separating out the five motifs that could possibly produce artifacts (orange numbers).

Yeast rRNA modifications have been extensively studied and well characterized ^17^. Using ELIGOS we identified 315 loci in yeast rRNAs (25S, 18S, 5S and 5.5S combined) with differential %ESB values. Of these, 67 loci matched known modified bases ^17^, covering 106 base positions of the total of 111 described modified bases (95%) which is a statistically significant finding, p-value of 7.2e^-84^. Our prediction did not capture five bases described to be modified (their regions did not produce %ESB elevated values; see Supplementary Figure S1). However, 248 additional loci were identified by ELIGOS that have not previously been described to undergo modification (Figure 2B). We checked for presence of the five triplets that were likely to produce artifactual results (*cf*. Figure 1F) and found that these represented 172 loci (54%). Interestingly, 35 of these have been previously documented as being methylated (Figure 2C). Thus, removal of these from the ELIGOS predictions would omit a number of experimentally verified modified base locations.

The data obtained with rRNA from *E. coli* were also compared to experimental documentation of *E. coli* rRNA base methylation ^18^. Of the 36 described methylated nucleosides described for the three bacterial rRNA molecules combined, our approach detected 33 (92%) with p-value of 1.3e^-28^ divided over 21 loci (Figure 2B). However, our data suggest that far more positions might contain modified bases. A total of 102 loci (42%) were due to the five triplets for which true and false signal could not be differentiated; 9 of these had been previously identified in the literature as being modified (Figure 1C). There were 3 previously described methylation sites that produced %ESB values lower than the cut-off threshold, or remained undetected due to presence of homopolymeric sequences (see Supplementary Figure S1).

The characterization of enzymes responsible for rRNA methylation in human cells is currently still incomplete ^19^. We compared our data with the Ribo-Methyl-seq data collected by Erales and colleagues ^20^ which at the time of analysis listed 106 2-O-methylation sites for rRNA of HeLa cells. Of the 413 loci predicted by ELIGOS, 58 overlapped with 79 positions of O-methylation sites (Figure 2B). Thus, 74% with p-value of 1.5e^-37^ of the data collected in RiboMethyl-seq were captured in our predictions. In a second analysis we compared our data to 3-dimensional human ribosome structural data derived from cryo-electron microscopy which can be employed to locate putative rRNA methylation sites with high confidence ^21^. The ELIGOS predictions captured around 78% with p-value of 5.1e^-83^ of those specific methylation sites. Interestingly, 35 of the 2-O-methylation bases reported by Erales *et al.* ^20^ were not captured in the data by Natchair *et al.* ^21^, and for 55 positions the opposite applied. For only 31 loci did ELIGOS predictions overlap with both published datasets (Figure 2B). For 164 predicted loci the results were inconclusive as they represented the five triplets for which no reliable data could be obtained (Figure 2C).

In summary, we were able to capture many of the known base modifications in rRNAs in yeast, *E. coli*, and human cells, as well as predict putative novel modified bases in rRNA. These results show that the method can detect a variety of potentially different modified bases simultaneously in native RNA.

### Comparison of dcDNA-seq and dRNA-seq from yeast transcriptomes

We next compared poly-A mRNA isolated from yeast cells grown in minimum media supplemented with glucose, and from cells that had switched to ethanol as a carbon source. For each condition three experimental replicates were analysed. The differences in read characteristics obtained from dcDNA-seq and dRNA-seq for the two transcriptomes are summarized in **Figure 3**. The sequence yield obtained per hour on the ONT flow cells (Figure 3A) was higher for dcDNA than for dRNA, due to the different motor proteins that control the rate of molecules passing through the nanopores (450 bases per second (b/s) for DNA and 80 b/s for RNA sequencing). The average % identities of both dcDNA and dRNA reads were comparable, around 88% (violin plot, Figure 3A). The base-calling step using Albacore software automatically classifies reads to fail or pass a specific cut-off. As seen in Figure 3B, on average 85% of the total dRNA reads but only 50% of dcDNA reads passed the default threshold of 7. The length of all reads combined (passed plus failed) indicated that the dcDNA reads were slightly longer than the obtained dRNA reads (Figure 3C).

**Figure 3.**
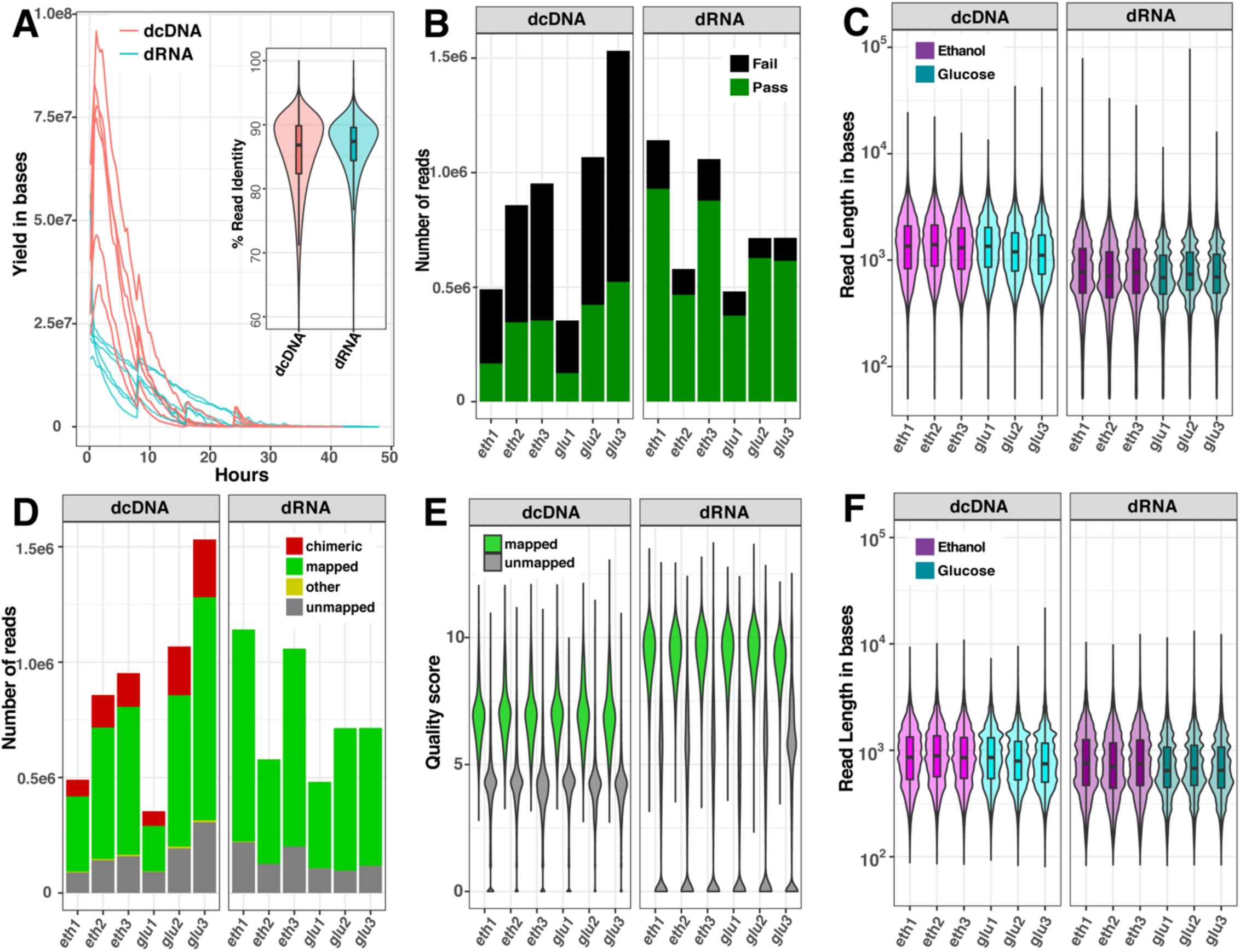
Comparison of read characteristics for six datasets of yeast RNA sequenced as dcDNA or dRNA. **(A)** Sequence yields per hour and violin boxplot of %read identity; **(B)** numbers of reads that passed (green) or failed (black) the quality score of 7 by Albacore software; **(C)** read length distribution of all reads combined (passed plus failed); **(D)** numbers of all reads that could be mapped to a reference genome; **(E)** quality score distribution of mapped and unmapped reads, and **(F)** read length distribution of the reads after removal of chimeric sequences. Data are shown for glucose-grown cells (glu) and for glucose-deprived cells (eth).

To explain the surprisingly high fraction of failed reads obtained with dcDNA, we re-evaluated the quality of total reads (passed plus failed) by aligning both dcDNA and dRNA reads onto a reference genome. As presented in Figure 3D, between 61% and 67% of the dcDNA reads could be mapped, while between 80 and 86% of the dRNA reads mapped to the reference genome. Of note was the relatively high fraction of chimeras in dcDNA (between 15 and 20%), while the fraction of unmapped reads (approximately 15%) did not significantly differ (p-value >0.05) between dcDNA and dRNA sequences. Further, the read quality score distribution of total reads differed between dcDNA and dRNA reads (Figure 3E), with higher scores for obtained for dRNA reads. Therefore, for the dRNA reads the recommended default of 7 was applied, while for dcDNA reads a less strict boundary quality score of 5 was deemed more suitable as transcript reads have a relatively shorter length than genomic DNA reads. This is in agreement with previous observations that shorter reads generated by ONT tend to produce lower quality scores ^22^. When the read length distribution was compared after removal of chimeric sequences from the dcDNA reads, this resulted in a comparable read length distribution for both sequencing strategies (Figure 3F).

The read counts of individual transcripts derived from the two different templates (DNA and RNA) were compared by scatter plot and a correlation matrix was constructed (Figure 4A). Within the same template, replicate experiments produced satisfying correlation coefficients (*r* =0.96 on average, range: 0.94-0.98), while on average an *r* of 0.92 (range: 0.90-0.94) was obtained when dcDNA and dRNA sequences were compared for the same growth conduction. We have recently demonstrated that the negative binomial statistic is a valid approach to analyze dRNA-seq data ^6^; here we applied that method to compare the adjusted p-values and the observed mean log2fold changes, as illustrated in Figures 4B and 4C, respectively. Even though the sequencing depth across the biological replicates varied, the results of both sequencing methods strongly correlated for transcriptomes that were obtained from cells grown under the same condition. Furthermore, biological functional enrichment was analyzed using Gene Ontology (GO) based on the dcDNA-seq and dRNA-seq data; the results were found to be highly consistent, as 332 GO-terms were identified in both datasets, and only 48 and 40 GO-terms were uniquely present in dcDNA-seq and dRNA-seq data, respectively (Figure 4D). The previously published conclusions on differential gene expression between the two compared culture conditions ^6^ did not change for the transcriptome sequencing data obtained here.

**Figure 4.**
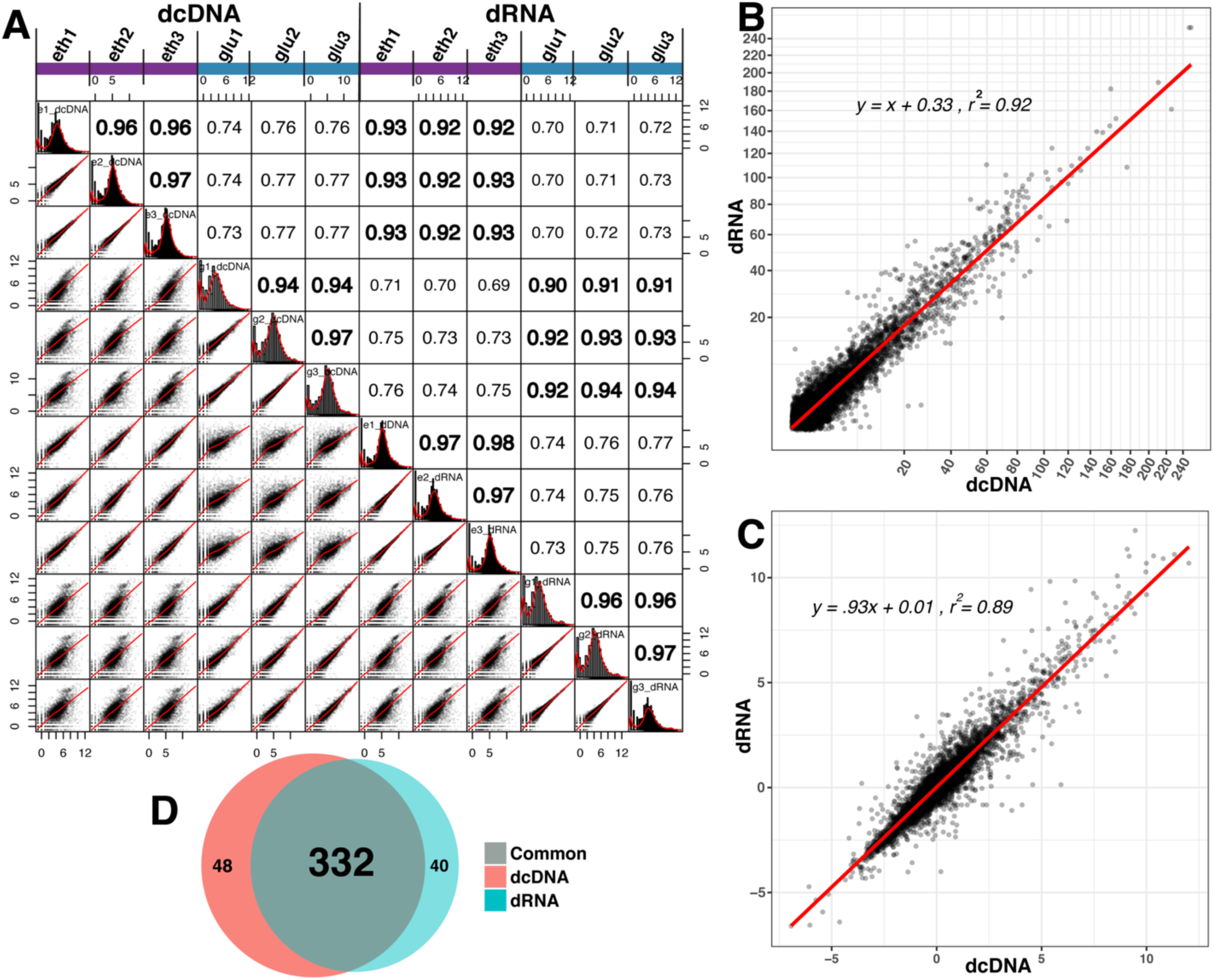
Comparison of transcript abundances based on dcDNA-Seq and dRNA-seq. **(A)** a combined scatter plot and correlation matrix. **(B,C)** Scatter plots showing the correlation of statistical values between all individual transcripts combined as identified by dcDNA and dRNA based on adjusted p-values **(B)** and on observed mean log2fold changes **(C)** derived from three biological replicates. **(D)** Venn diagram of GO-terms identified in dcDNA and dRNA datasets.

### Over-representation of the artifactual triplets in modified base predictions

ELIGOS predictions were next applied to the yeast transcriptomic data described above, complemented with a third dataset of mRNA isolated from *S. cerevisiae* strain DBY746 grown in rich media (YPD) ^5^. A fourth dataset was added which consisted of mRNA isolated from human lymphoblastoid cell line, GM12878, which is part of the publicly available Oxford Nanopore Human Reference Dataset. Using the same statistical cut-off as defined in the previous section, approximately 18,000 positions in the yeast datasets and 85,000 positions in the human cell line data were identified with differential %ESB positions. Comparing the four bases, the highest fraction of differential %ESB positions in all four datasets combined captured by ELIGOS was for cytidine, comprising 40% of the total differential %ESB positions on average (see Supplementary Table S2). We evaluated enrichment of motifs surrounding the differential %ESB positions and found four motifs that were consistently overrepresented in all four datasets, as illustrated in Figure 5A. The overrepresentation was strongest for motif U<u>C</u>U (with the underlined C being the identified base). The motif uc<u>U</u>CC (with variants UCC<u>U</u>C and C<u>U</u>CC for yeast strain DBY746 and human RNA, respectively) was overrepresented for positions containing uridine, and C<u>A</u>C (UC<u>A</u>C in human RNA) and C<u>A</u>UG (with variants u<u>A</u>uGG and C<u>A</u>uGG) for those containing adenine. Of note is that these motifs all contained the five over-represented triplets that had been identified as producing unreliable findings by the IVT luciferase analysis.

**Figure 5.**
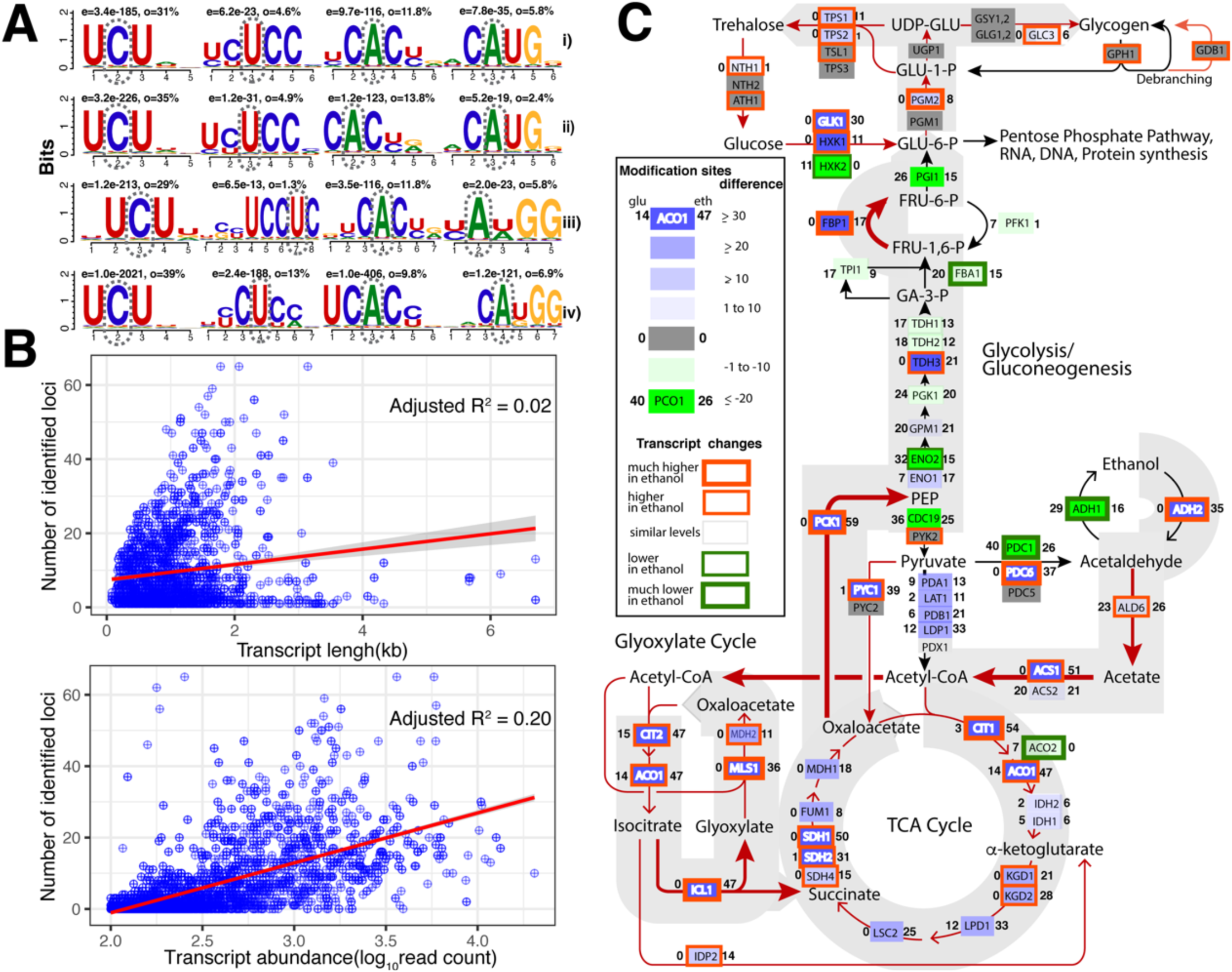
Capturing RNA modification and structural signatures inferred by ELIGOS in 4 datasets of mRNA. **(A)** Logo plots of the most common motifs around the differential ESB positions identified by ELIGOS (indicated by the dashed line ovals) in the transcriptomes from yeast strain CEN.PK113-7D grown in glucose (i) and in ethanol (ii), yeast strain DBY746 grown in YPD (iii) and from a human cell line (iv), for (left to right) cytidine, uridine or adenine. Above each plot, **e** refers to the e-value of the motif, and **o** reports the occurrence of the motif. **(B)** Scatter plots of the yeast data sets with linear regression lines, showing no correlation between transcript length (top) and weak correlation between transcript abundance (bottom) and their number of identified inferred RNA modification loci. (**C**) Concerted analysis of differential gene expression and RNA modifications as inferred by ELIGOS on the central metabolic pathway during the diauxic shift of yeast. The green and blue boxes represent the difference in number of inferred RNA modifications in individual transcripts that are higher in glucose and ethanol, respectively, with the numbers of inferred RNA modifications on the left and right of the boxes, respectively. The grey boxes represent transcripts that have no inferred RNA modifications detected. The edges represent the fold changes of transcript abundances.

The identified differential %ESB positions were cleaned for the four motifs for which artifactual and real signals could not be distinguished, resulting in a ∼57% reduction (see Supplementary Table S2). This retained 8,889 differential %ESB loci in the mRNA dataset of yeast grown on minimal medium with glucose, corresponding to 691 transcripts. Likewise, 6,806, 5,488 and 24,702 differential ESB loci were identified in yeast using ethanol, yeast cultured in YPD and in the human cell line dataset, corresponding to 788, 758 and 3,234 transcripts respectively (only canonical transcripts were considered, excluding isoforms).

### Association of inferred RNA modifications with transcript abundance and length as exemplified by yeast

We next evaluated whether an association exists between transcript abundance or transcript length and their number of inferred RNA modification loci, per dataset. No strong correlation was found between the number of differential ESB loci and transcript length, in all four datasets (R^2^ <0.0005 for yeast on glucose, <0.007 for yeast on ethanol, <0.007 for yeast in YPD and 0.01 for human cell transcripts, respectively; see Supplementary Figure S3). The analysis of the three yeast datasets combined is shown at the top of Figure 5B.

A weak linear trend was observed between highly abundant transcripts (covered by ≥100 reads) and their number of differential ESB loci (Figure 5B, bottom). This weak positive correlation was found in all four datasets (R^2^ = 0.20 for yeast on glucose minimal media, 0.17 for yeast on ethanol minimal media, 0.35 for yeast in YPD and 0.12 for human cell transcripts, respectively; see Supplementary Figure S3). Lack of a correlation between inferred RNA modification status and expression levels can be exemplified by zooming in at some of the hyper-modified transcripts, defined as having >20 differential ESB loci, in the yeast datasets. These covered 104, 100, and 56 transcripts from cells grown on glucose, ethanol, and YPD, respectively (Supplementary Figure S4 illustrates the overlap between these datasets in a violin jitter plot and an Upset plot and more details of individual gene is provided in Table S4). Some of the hypermodified transcripts were extremely abundant during growth on ethanol, e.g., carnitine acetyltransferase (YAT1, with >5600 reads) and the chromosomal gene for Hexose Transporter Induced by Decreased Growth (HXT5, >3600 reads), but the transcript of the Shmoo tip protein (HBT1) was much less abundant (∼250 reads), while these three transcripts all contained 65 modification sites.

A correlation between base modification levels and transcript abundance was obvious, however, when zooming in at specific pathways. This is exemplified by the central metabolic pathway shown in Figure 5C. We mapped relevant transcripts and their number of inferred RNA modification loci to simultaneously assess the effect of transcriptional and posttranscriptional regulation during metabolic reprogramming required for the diauxic shift. The presented global overview shows the well-known adaptations ^23^ of yeast cells as they switch from glucose to ethanol, by changing gene expression of a number of key enzymes. In addition to transcriptional regulation, we found many transcripts that had undergone changes in base modifications under these conditions. Examples are genes under regulation to switch from glycolysis to ethanol utilization (ADH2 and ACS1), key genes regulating the TCA cycle activity (CIT1, ACO1 and SDH1,2), the glyoxylate shunt (ICL and MLS1) and the key enzyme in gluconeogenesis (PCK1). On the other hand, the enzymes involve in glycogen-trehalose homeostasis were transcriptionally regulated while hypo-modified (e.g., NTH1, TPS1,2, GLC3, PGM2) or not modified (e.g., ATH1, TSL1, GPH1, GDB1). Interestingly, acetaldehyde dehydrogenase ALD6 was upregulated when cells utilized ethanol but its transcript modification only marginally differed between the conditions. These results indicate there exists a complex association between transcript modifications and metabolic reprogramming.

### The Human Transcriptome: Capturing known m6A and RNA G-quadruplexes

Lastly, we analyzed the transcriptome of the human cell line and examined the two most abundant motifs surrounding the modification sites captured by ELIGOS, shown in Figure 6. (The most abundant identified motifs of all four datasets is shown in Supplementary Figure S5.) Interestingly, the two most abundant motifs in the human dataset both have known biological relevance (Figure 6A, B). The first motif GG<u>A</u>CH (Figure 6B) is the known DRACH consensus sequence for m6A recognition sites, where D = A/G/U, R = A/G, and H = A/C/U ^24, 25^. This motif is recognized by epigenetic ‘reader’ proteins (YTH RNA-binding domain proteins ^26, 27^). YTH RNA-binding domain proteins control several important pathways, including neural development in humans ^28^. The motif in Figure 6A represents the most abundant base methylation site identified to date, and is the best studied case of 6mA RNA methylation in eukaryotes. We identified this as the most abundant adenine motif with of e-value of 5.1e^-224^ and 14 % occurrence, corresponding to 965 transcripts (see Supplementary Table S4 for details on numbers of loci/motifs in each transcript). For these 965 transcripts, we analyzed the positions of the identified DRACH motifs along each transcript and compared this to the sequencing depth of dRNA-seq over the location of the transcripts; the data are presented in a standardized coordinate plot in the lower part of Figure 6A. This identified a clear preference for the DRACH motif to be present at the gene-bordering flank of the 3’ untranslated region (UTR), which agrees with previous studies ^24, 29, 30, 31^. The second motif (Figure 6B) represents the most abundant guanine motif with e-value 6.1e^-89^ and 41% occurrence, corresponding to 1250 transcripts (see Supplementary Table S4 for details). This motif G<u>G</u>AGG was identified to form RNA G-quadruplexes (rG4s) ^32^. By plotting the standardized coordinates of the location of this rG4s motif and comparing it to the sequencing depth of dRNA-seq (Figure 6B, lower panel), we found an even distribution of the motif with a small bias for the gene-bordering flank of the 3’ untranslated region (UTR).

**Figure 6.**
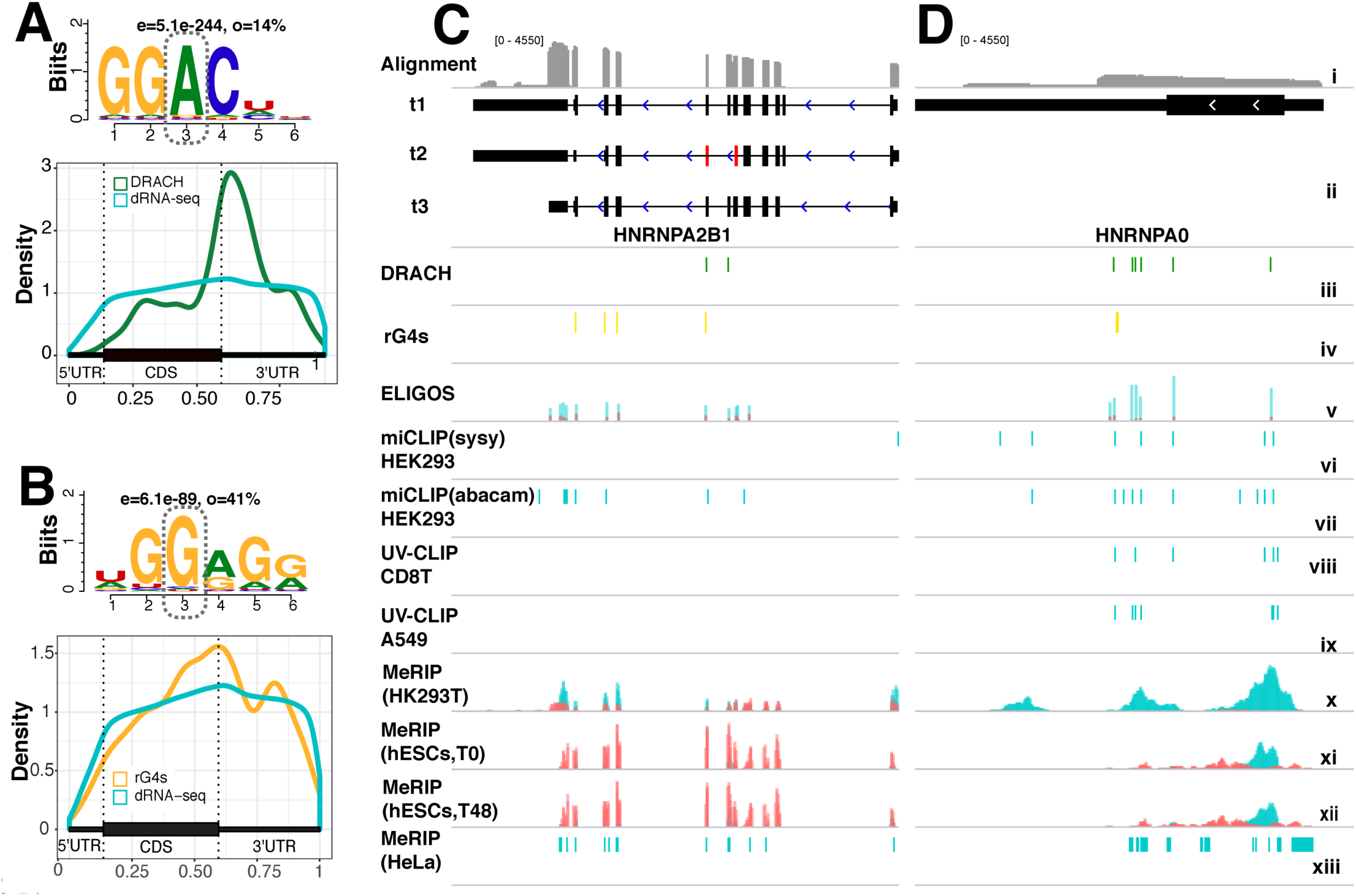
Epitranscriptome of human cell line CEPH1463s. **(A)** Logo plot of the DRACH motif surrounding m6A identified by ELIGOS, with below it the standardized coordinate plot of 995 transcripts containing the motif to illustrate its preferential position in 3’ untranslated regions. **(B)** Logo plot of the RNA G-quadruplexes (rG4s) motif with below it the standardized coordinate plot of the 1250 transcripts containing the motif. Other motifs identified in the yeast datasets are shown in Supplementary Figure S6. **(C, D)** Examples of selected transcripts hnRNP A2/B1 **(C)** and hnRNP A0 **(D)** in which both the DRACH and the rG4s motifs were found to be modified. A comparison is shown in IGV Genome Browser of our predictions and previous studies conducted with different human cells and different m6A profiling methods. The tracks show (from top down): *i)* alignment coverage depth of dRNA reads of the transcripts; *ii)* isoform architecture showing (D) transcripts t1, t2 (missing exons 7 and 8, shown in red), and t3; *iii)* location of ELIGOS identified DRACH motifs (green); *iv)* location of ELIGOS identified rG4s motifs (yellow); *v)* %ESB of dRNA (cyan) and dcDNA (red) sequences at the differential %ESB loci for adenine as identified by ELIGOS; *vi)* m6A individual-nucleotide resolution crosslinking and immunoprecipitation (miCLIP) data of HEK293 cells using SySy m6A antibody enrichment ^24^; *vii)* miCLIP data of HEK293 cells using Abacam m6A antibody enrichment ^24^; *viii)* UV crosslinking and immunoprecipitation (UV-CLIP) data of CD8T cells ^29^; *ix)* UV-CLIP data of A549 cells ^29^ *x)* methyl-RNA immunoprecipitation (MeRIP) peak data of HEK293T cells ^30^; *xi)* MeRIP peak data of hESCs cells at time point T0 ^52^; *xii)* MeRIP peak data of hESCs cells at time point T48 ^52^. All MeRIP peak data were plotted based on the read coverage depth of 6mA enriched (cyan) and the reference sequencing library (red); *xiii)* MeRIP peak region data of HeLa cells ^35^. A zoomed output is shown in Supplementary Figure S6.

Presence of both the DRACH and rG4s motifs in a single transcript may imply complex post-transcriptional regulation. To give an example, the transcript of RNA binding protein hnRNP A2/B1 (which promotes primary microRNA processing, is involved in splicing regulation and potentially serves as a m6A reader ^33^), can itself undergo alternative splicing to produce two experimentally confirmed isoforms and another rare isoform associated with presence or absence of exons 1, 7 and 8 ^34^. In the transcripts of this gene we identified 2 DRACH and 4 rG4 motifs containing modified bases, including one of each in exon 7 and a DRACH motif in exon 8 (Figure 6C). Interestingly, ELIGOS identified other %ESB loci where DRACH motifs were absent that have been described as containing m6A, detected in miCLIP(abacam) data of HEK293 cells ^24^, in MeRIP data of HK239T^30^ and in MeRIP data of HeLa cells ^35^. The inconsistency of m6A detection across different studies indicates highly complex and dynamic cellular regulation of methylation patterns that is cell type specific. The coverage plot from the alignment of dRNA-seq reads indicates that the third isoform with the shortest 3’ UTR was the most abundant isoform of hnRNP A2/B1 in the investigated transcriptome, while minor amounts of the first isoform were also detected, indicated by the low coverage depth of the first exon. The abundance of the second isoform, which produces the shortest protein among the three isoforms (lacking exons 7 and 8), was too low to be detected. This shortest isoform lacks a glycine-rich region and other important domains and posttranslational modification sites necessary for protein function. Therefore, inclusion of exons 7 and 8 is important for protein function, and the presence of both the m6A and rG4s motifs, containing modified bases as predicted by ELIGOS, is most likely involved in this inclusion to promote translation of the biologically active isoform. A role of base modification in these motifs involved in their biological functions can be assumed, in line with studies that have shown that exon inclusion into mRNAs is promoted by m6A through YTHDC1 ^36^ and by secondary structures formed by rG4s ^37^.

A second example of a transcript containing both DRACH and rG4s motifs is hnRNP A0, heterogeneous nuclear ribonucleoprotein A0 that contains six and one of these, respectively (Figure 6D). ELIGOS predictions highly agreed with all experimental miCLIP data, even at single nucleotide resolution (see Supplementary Figure S6 of a zoomed view), and with MeRIP studies on the region that has high depth coverage of dRNA-seq. In addition, the differential %ESB of adenine in this transcript that was filtered out by the artifactual triplet CAC was detected by miCLIP(SySy) ^24^ as an m6A modification. This observation again supports the undistinguishable RNA modification from artifactual signals (see Supplementary Figure S6).

### Discussion

The major fraction of sequencing errors by ONT, which captures single molecule sequences, is derived from stochastic noise that can be corrected for by consensus base calling from reads pileup ^38^. The consensus error correction approach typically results in correction of sequencing errors when DNA is sequenced, however ∼1% of the total errors typically need to be further polished by short reads ^38^. Sequencing of native RNA results in more errors, as we found higher %ESB scores for this template (Figure 1A). We demonstrated that this is a combined effect of ribonucleoside modifications as well as presence of secondary structures. The ONT technology is still in its infancy and especially base calling software for RNA is not as well developed yet as for DNA; for example, the RNN model used for RNA has only been updated once so far, while the DNA model is more advanced ^15^. Our observations that five particular triplets are overrepresented in high %ESB scores (Figure 1F) can assist in further fine tuning the base calling software in the near future, which we expect will improve the base calling model for RNA.

When present, base modifications and secondary structures of nucleic acids alter the ionic current signal recorded during ONT sequencing, leading to errors that are inherent to the application of helicase and pore protein for pore passage. We developed ELIGOS for determining a comparative error analysis of long read sequences, as this can be used as a signature to recognize base modifications and secondary structures. By sequencing *in vitro* transcribed RNA, we are able to compare the errors recorded with modified RNA with that of naked RNA or cDNA signals. Although similar results were obtained (Figure 1E), the use of dRNA sequences from naked RNA obtained by IVT as the reference is more suitable to eliminate the systematic errors caused by particular triplets as well as secondary structures. Nevertheless, construction of *in vitro* transcripts to study genome-wide RNA modifications is not trivial, and the use of cDNA as a reference results in proper identification of secondary structures such as those caused by the rG4 motif (Figure 6). This capability can be potentially extended to study RNA secondary structures.

Distinct error signatures were identified by ELIGOS between native, modified RNA and cDNA templates at base resolution, which captured most of the known RNA methylation sites, for all four bases simultaneously, despite inherent differences in methylation of these bases. This was demonstrated in yeast, *E. coli* and human RNA. This provides a promising approach to detect expected as well as novel RNA methylations and base modifications directly from native RNA sequences. This capability is superior to traditional methods that can detect one type of methylation at the time only and require complex experimental procedures. Moreover, based on the same principle, ELIGOS can be applied to identify DNA modification by the comparison of the errors between native DNA and cDNA or a PCR product as shown in the Supplementary Figure S7. This potential will need to be further investigated and compared with existing methods for direct DNA modification detection using ONT ^39, 40^ or PacBio ^41^ sequencing.

The procedure can result in possible high false positives from artifactual signals, as was demonstrated for five triplets that caused errors in the nanopore sequencing signals that were irrespective of presence of 5moU in the IVT experiment. Such systematic errors can be filtered out from the ELIGOS results if different mRNA datasets can be compared, helping to reduce false positives, at the cost of removing true signals that can be presented by these sequences. Using this approach, we were able to uncover known biologically relevant motifs containing m6A RNA methylation and rG4 secondary structures. ELIGOS can specifically identify the location of RNA modifications but it cannot tell the exact type of RNA methylation. This is a limitation of the approach and it would require further investigations to determine the nature of the RNA modification loci inferred by ELIGOS by using such traditional technique of LC–MS/MS approach ^14^. This will be a complementary approaches for epitranscripome profiling.

Systemic analysis of transcriptional and epitranscriptional regulations would provide a better understanding of cellular adaptions. We applied our method here to either rRNA or poly-A RNA transcripts. It has previously been reported that in a given cell population, even rRNA methylation patterns can be heterogeneous ^42^ whose nature may depend on dynamic processes taking place at a cellular level, and on the stage and cell type that can be used as a marker for cancer ^42^. We have further demonstrated (Figure 5) that metabolic reprogramming of the central metabolic pathways of yeast during the diauxic shift from glycolysis and alcoholic fermentation to aerobic respiration and gluconeogenesis relied on regulation of both transcript abundance and base modifications. To our knowledge this has not been previously reported in the literature. This kind of regulation coupling was also found in RNA undergoing methylation-mediated pathways in cancer cells, so that our method now opens a new strategy to study carcinogenesis ^43^.

The limitations of our method is that for a number of sequence triplets, false-positive signal could not be distinguished from real signals. Moreover, the method identifies the location of putative modification sites but not its nature, whose identity would need further investigations. Besides, the input data for our method depend on the results obtained from base calling and long read aligner software as a prerequisite, therefore the accuracy of these steps will influence the final result. Lastly, it is possible that the method is over-reporting the number of predicted modified bases due to the noisy nature of ONT output. Nevertheless, this systematic sequence approach to determine the epitranscriptome of a cell can be used to direct an experimental work flow, especially since expression levels can simultaneously be considered.

In conclusion, this study provides a concrete foundation to study native RNA sequences that carry important information on RNA modifications, secondary structures and possible other features responsible for sequence errors. Detailed investigations to dissect the complex properties of RNA from detected error signatures is now feasible. Our ELIGOS software is publicly available and can be used to detect possible RNA modification sites and secondary structures quickly, on a global transcriptomic scale. Moreover, ELIGOS can be used as a diagnostics tool to improve the base calling algorithm of nanopore sequencing. We envisage that sequencing of native RNA will become a powerful and versatile tool to advance RNA biology.

## Methods

### *In vitro* transcription of luciferase mRNA

*In vitro* transcription (IVT) to produce mRNA of the luciferase gene (L-7602 CleanCap™ Firefly Luciferase, TriLink Biotechnologies, San Diego, CA, USA) was carried out with standard ribonucleotides and with incorporation of 5-methoxyuridine (5moU, TriLink Biotechnology). The produced mRNA containing a poly-A tail was purified using AMPureXP beads (Beckman Coulter, Brea, CA, USA) and eluted using nuclease-free water.

### Culture conditions and RNA extraction

Yeast RNA used for ribosomal RNA was isolated from *S. cerevisiae* strain S288C grown overnight at 30°C in 15 mL medium containing 10 g/L yeast extract, 20 g/L peptone, and 20 g/L glucose. RNA was extracted using the ZymoBIOMICS Quick-RNA Fungal/Bacterial kit (Zymo Research, Irvine, CA, USA) according to the manufacturer’s protocol. The yeast poly-A RNA used to compare the transcriptome of different culture conditions is the same as previously described ^2, 6^. *S. cerevisiae* strain CEN.PK113-7D was cultivated overnight in minimal medium containing 20 g/L glucose as the carbon source. Cells were harvested during mid-exponential growth on glucose and during late-phase growth, when the cells had switched to aerobic respiration and consumed ethanol due to glucose limitation. The same RNA aliquots were used to produce dcDNA sequences as described below. The data from three independent replicate experiments were used, producing 12 sequence data sets (three each for dcDNA-seq and dRNA-seq from either glucose-grown (glu) or glucose-depleted cells (eth)).

*Escherichia coli* strain ATCC 11775 was cultured overnight at 37°C in 25 mL of Luria broth (LB) and following centrifugation the cell pellet was resuspended in 250 µL, to which 750 µL of TRIzol reagent (Life Technologies, Carlsbad, CA, USA) was added. Following incubation for 5 minutes at room temperature, 200 µL of chloroform were added. Phases were mixed by inverting the tube 15 times and then incubated for 10 min. Following centrifugation at 12,000 x g for 5 min at 4°C, 400 µL of the aqueous phase was removed and the RNA it contained was cleaned using the Direct Zol kit (Zymo Research).

Human cell line KTC-1 (human papillary thyroid cancer cell line) was grown to 85-90% confluence in 10cm dishes in RPMI media supplemented with 10% fetal bovine serum utilizing standard techniques. The cells were rinsed twice with cold, sterile PBS after which 700 µl TRIzol reagent (Life Technologies) was added. Following incubation for 5 min at room temperature, the cells were collected and mixed with 700 µl absolute ethanol. RNA isolation was performed with the Direct-Zol RNA mini prep Kit (Zymo Research) as per manufacturer’s instructions. Total RNA was eluted in 20µl RNase/DNase free water and stored at -80°C. As most RNA in these samples represented ribosomal RNA, the template was completely sequenced to obtain rRNA reads.

The total RNAs for the rRNA experiments were firstly add poly-A using *E. coli* Poly(A) Polymerase (New England Biolab, UK), following the manufacturer’s protocol, then used for sequencing library preparation.

### Library preparation, dcDNA-Seq and dRNA-Seq by ONT

A total of 530∼600 ng total yeast RNA was enriched for poly-A RNA by means of oligo(dT) beads and this was used to prepare both libraries. The dcDNA library was produced using the SQK-DCS108 kit (ONT, Oxford, UK) which includes an RT step but no amplification step. RNA was then converted to double strand DNA, after which ligation of the adaptor attached the motor protein (Supplementary Figure S8). The library was loaded directly onto a flow cell for sequencing using a MinION Mk1B. Preparation of the library for dRNA-seq, SQK-RNA001 was used, only required an RNA stabilization step by formation of DNA-RNA hybrids through reverse transcription. After this, the motor protein was attached to the RNA strands specifically. Each library was loaded onto a flow cell for a 48 hours sequencing run lasting. Direct sequencing of the poly-A RNA (dRNA) was performed on a single R9.5/FLO-MIN107 flow cell.

### Bioinformatics and statistical analysis

#### Data processing and mapping of reads

The ONT raw data (.fast5 files) generated by MinKnow software (version 1.7.14) were converted to basecalled .fastq files using the local-based software Albacore version 2.1.3. This step automatically classifies failed and passed reads based on a specific cut-off for mean quality scores of 7 and only reads >200 bases were included. The ONT reads in standard fastq format were aligned to the reference sequences using Minimap2 to generate a BAM file. The dRNA reads were converted to DNA sequences and reverse complement sequences of dcDNA reads were generated before alignments. For analysis of mapping results of yeast, we employed SAMtools (version 1.6) to investigate the BAM files and to classify sequence reads into categories of mapped, unmapped, chimeric and other reads based on standard CIGAR string information.

#### Comparative errors analysis and development of ELIGOS software

The ELIGOS software was developed to compare the error signals between dRNA and dcDNA/cDNA sequences. The percentage of errors at a specific base (%ESB) is defined as the percentage of the sum of substitutions, insertions and deletions of individual positions over total mapped reads obtained from read alignment results based on the reference sequence. Each pair of BAM files, together with reference sequences and transcript annotation files in bed12 format, was used as the input of the ELIGOS software. The calculations of %ESB through the pysam module and the statistical tests (explained below) by R were performed using individual base positions of transcripts over the reference sequences with multithread parallelization architecture. The software was then applied to the rRNA and the mRNA sequencing datasets. ELIGOS is written in python and is available at https://bitbucket.org/piroonj/eligos.git. The difference of the %ESB between dRNA and dcDNA sequences of identical positions in the reference sequences were evaluated using either Fisher’s exact test for a single 2×2 consistency table (one biological replicate) or Cochran–Mantel–Haenszel test for multiple (more than one biological replicate) 2×2 consistency tables of independence. The statistical p-values were further adjusted for multiple testing using the Benjamini-Hogberg method. The adjusted p-values <1e^-50^ and odds ratios (errors presented in dRNA sequence over errors presented in dcDNA sequence) ≥2 were used as cut-offs to reject the null hypothesis that the errors at the individual base of dRNA and dDNA sequences were equal. Furthermore, a cut-off of ≥25% ESB in dRNA sequence was used as additional filter to remove noise due to the error-prone long reads as illustrated in Figure 1A. Some interesting regions were explored at the signal-level through the re-squiggle signal approach using Tombo software version 1.4 (https://github.com/nanoporetech/tombo.git).

For ribosomal RNA investigations, the fastq files were aligned onto a reference genome sequence (for *S. cerevisiae*: NR_132209.1, NR_132215.1, NR_132213.1, and NR_132211.1 combined; for *E. coli*: positions 232785-23568, 1046691-1048228 and 232576-232686 from NZ_KK583188.1; and for *H. sapiens* NR_023363.1, NR_003287.4, NR_146119.1 and NR_145819.1 combined) using minimap2 software ^44^ to obtain BAM files of the sequences.

#### Evaluation of mRNA sequencing characteristics

The yeast dRNA reads from strain CEN.PK113-7D were downloaded from the SRA database (accession number SRP116559), and after generation from the same sample aliquots, the corresponding dcDNA reads. The sequence reads from yeast strain DBY746 grown in YPD were downloaded from BioSample SAMN07688322 ^5^. A fourth dataset was added which consisted of mRNA isolated from human cell line, GM12878, which is part of the publicly available Oxford Nanopore Human Reference Dataset (https://github.com/nanopore-wgs-consortium/NA12878/blob/master/RNA.md) under creative license 4.0 ^8^. All data generated in this study were deposited in the SRA database (accession number SRP166020).

#### Differential gene expression evaluation

We followed the workflow to analyze differential gene expression of yeast transcripts as previously described previously ^6^. In brief, the read count table of individual transcripts for the dcDNA and dRNA sequences were generated using Bedtools version 2 ^45^. We then employed the DESeq2 package ^46^ to calculate adjusted p-values of individual transcripts between the two compared growth conditions. Consequently, functional gene enrichment analysis based on GO annotation was performed using the PIANO package ^47^.

#### De novo motif discovery

The sequences of 20 bases surrounding the differential %ESB of all A, T, C, or G positions identified by ELIGOS were extracted based on the reference sequence and these four separate datasets were analyzed using XXmotif software ^48^ to identify conserved motifs. The selected results of common motifs across the four experimental datasets are illustrated as logo plots with e-values and percent occurrence.

#### Genomic locations of loci and transcripts comparison

The relative location of considered loci with reference to gene position was compared using Bedtools version 2 ^45^ and the GenomicRanges package ^49^. The results were summarized in Venn diagrams using ChIPpeakAnno ^50^ or Upset plots using UpsetR ^51^.

#### Statistical analysis

Fisher’s exact test was used for a single 2×2 consistency table (one biological replicate) and the Cochran–Mantel–Haenszel test for multiple (more than one biological replicate) 2×2 consistency tables of independence. The statistical p-values were further adjusted for multiple testing using the Benjamini-Hogberg method. These statistical tests were used to compare %ESB of individual bases. The results from Fisher’s exact test were used to generate Figures 1E, 2B, 2C, and the human cells dataset. Cochran–Mantel– Haenszel test was used for the yeast datasets. Negative binomial statistics of the DESeq package was employed for differential expression analysis of the yeast grown in minimal media and shown in Figure 4B. Statistical analysis of gene-set enrichment was performed under PIANO package and shown in Figures 4C, D. Student’s *t*-test was used in Figures 1A, 2A to compare populations of %ESB between dRNA and dcDNA. Wilcoxon signed-rank sum tests were employed to test the difference of means between two considered populations in Figure 1F, to compare %ESB between of the five artifactual triplets among dRNA^O^, dRNA^U^ and dcDNA. Statistical significance of reported comparisons between methylation predictions and published experimental results of rRNA were calculated using hypergeometric test to reject the null hypothesis that the findings were produced by random events. The statistical analyses were performed using the R suite software.

## Supporting information

## Acknowledgments

### General

We thank Rui Perira for providing the RNA material from our previous collaboration.

### Funding

This work was partly supported by the Helen Adams and Arkansas Research Alliance Endowed Chair, and the National Institute of General Medical Sciences of the National Institutes of Health (awards P20GM125503 and 1P20GM121293).

### Author contributions

IN designed and conceived the project. TW performed MinION sequencing for dRNA-Seq and dcDNA-Seq as well as data submission. PJ, IN performed computational analysis and together with TMW interpreted the data. DU, TMW, ATF, NSA and MLJ participated in the study design. IN, TMW, TW, PJ wrote and edited the manuscript. All authors have read and approved the final version.

### Competing interests

The authors declare no competing interests.

### Data and materials availability

All data generated in this study were deposited in the SRA database (accession number SRP166020). ELIGOS is available from https://bitbucket.org/piroonj/eligos.git.

