## Supplementary material for "Decoding the Epitranscriptional Landscape from Native RNA Sequences"

### Supplementary Information Figures S1-S7

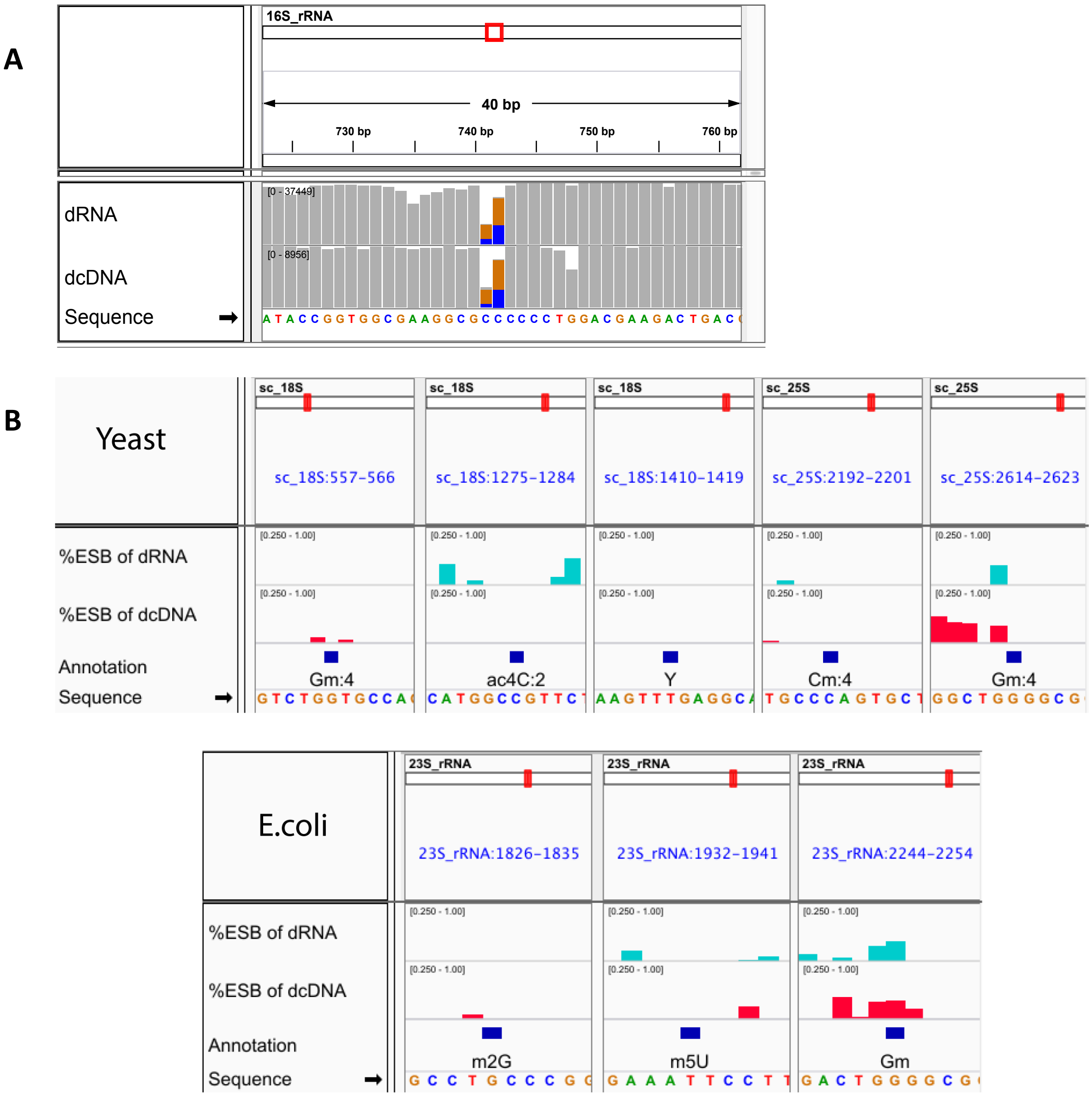


**Supplementary Figure S1.**

**(A)** The effect of homopolymeric stretches on %ESB found in both dRNA and dcDNA, with $C_6$ as an example. The panel shows a snapshot of the IGV genome browser.

**(B)** Snapshots of IGV genome browser showing regions with known methylations of yeast and *E.coli* rRNA that were not captured by the %ESB comparison between dRNA and dcDNA using ELIGOS. The upper panel shows the 5 uncaptured rRNA methylations of yeast rRNA and the lower panel the 3 uncaptured rRNA methylation sites of *E. coli*. Most of these produced no %ESB signal with the native RNA template. The last column of both panels shows the strong impact of homopolymers present in the vicinity of a modified base that overruled any possible specific signal.

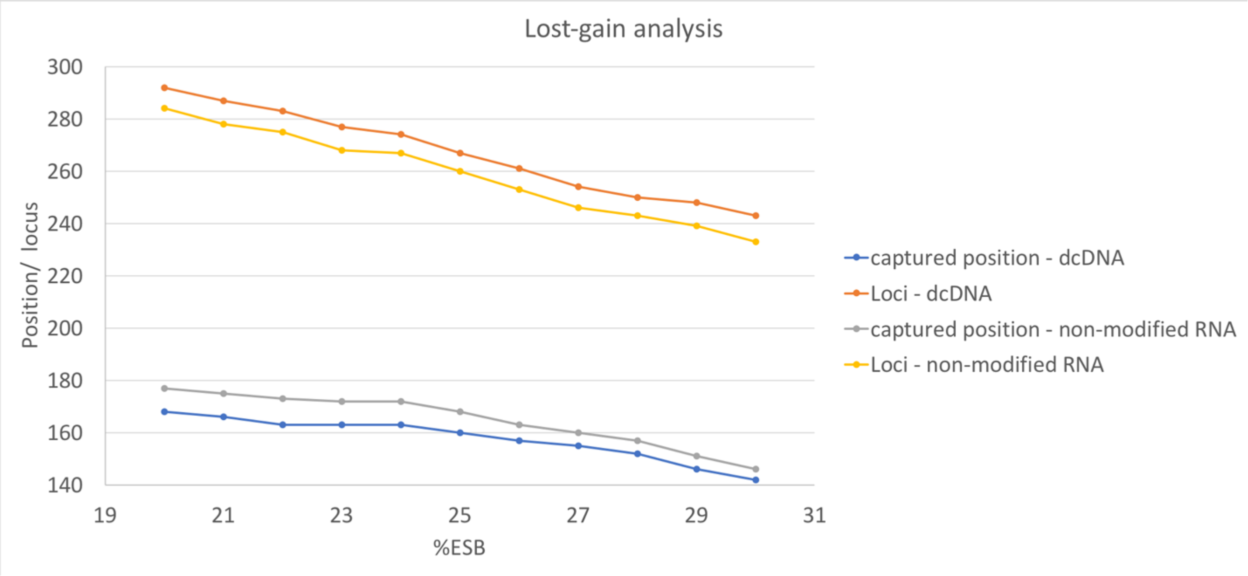


**Supplementary Figure S2**. Detailed loss-gain analysis to determine the optimal %ESB cut-off (range: 20-30%) at an odds ratio of ≥2 and adjusted p-values $<1e^{-50}$. The plot shows the number of identified differential loci (top two curves) and positions of 5moU (lower two curves) that were captured when either $dcDNA^{O}$ (blue, orange) or non-modified $dRNA^{U}$ (yellow, grey) was used as the reference. A plateau is visible between 22% - 24% for captured 5moU positions with both references, and between 23% and 24% for the total loci captured with $dcDNA^{O}$, which indicated the signal is noisy in this range. Beyond 24%, a more stringent %ESB results in increasing loss of total captured positions and loci. Based on this analysis we consider 25% (the first integer percentage beyond the observed plateau) as optimal.

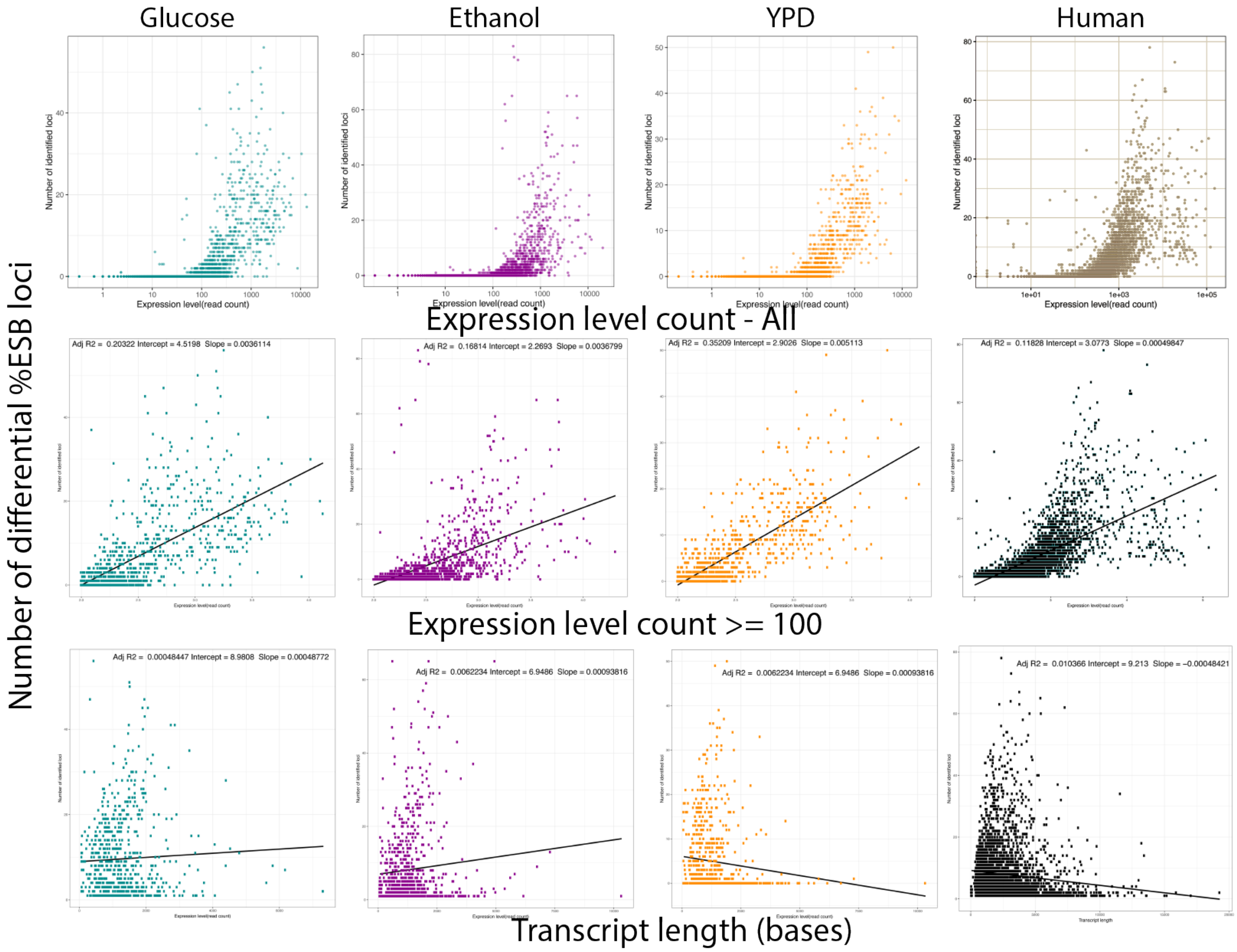


**Supplementary Figure S3.** Scatter plots of the four datasets, between number of identified differential loci of individual transcript (y-axis) with their expression level counts (x-axis), for all transcripts (top row), transcripts with expression levels ≥100 read counts (middle row) and for transcripts that contained at least one differential ESB locus (bottom row). The top panels show that the differential %ESB loci were identified in transcripts with expression levels from around 100 reads upwards. The summarized results of linear regressions are overlaid on each plot.

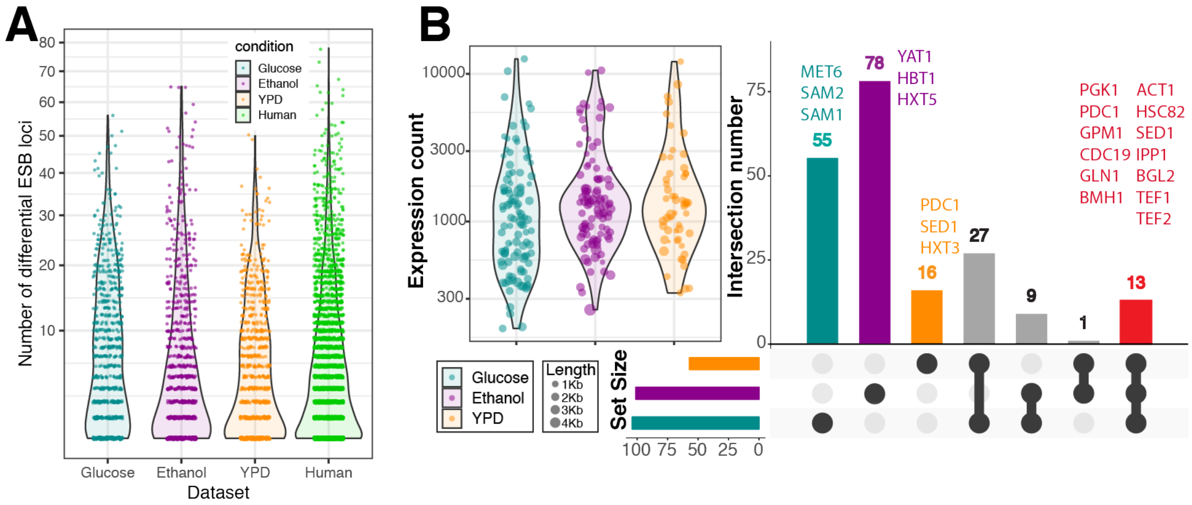


**Supplementary Figure S4. (A)** The distribution of the number of differential %ESB loci for individual transcripts (the genes lacking such loci were discarded from the plots), as identified by ELIGOS across the different four datasets. **(B)** Summary of hyper-modified transcripts in the yeast transcriptome. The violin scatter plots show the distribution of expression levels of hyper-modified transcripts with dot sizes representing the transcript length. The Upset plot shows the overlap between hyper-modified transcripts derived from the three yeast datasets. Dark grey dots at the bottom identify presence in a given dataset. The gene names of the 13 hyper-modified transcripts identified in all three yeast data sets are presented in red. The most strongly hyper-modified transcripts that were specific for each data set are presented by their gene name, for growth on glucose (purple), growth on ethanol (magenta), and in orange for strain S2988C grown in YPD, with pairwise interceptions in gray.

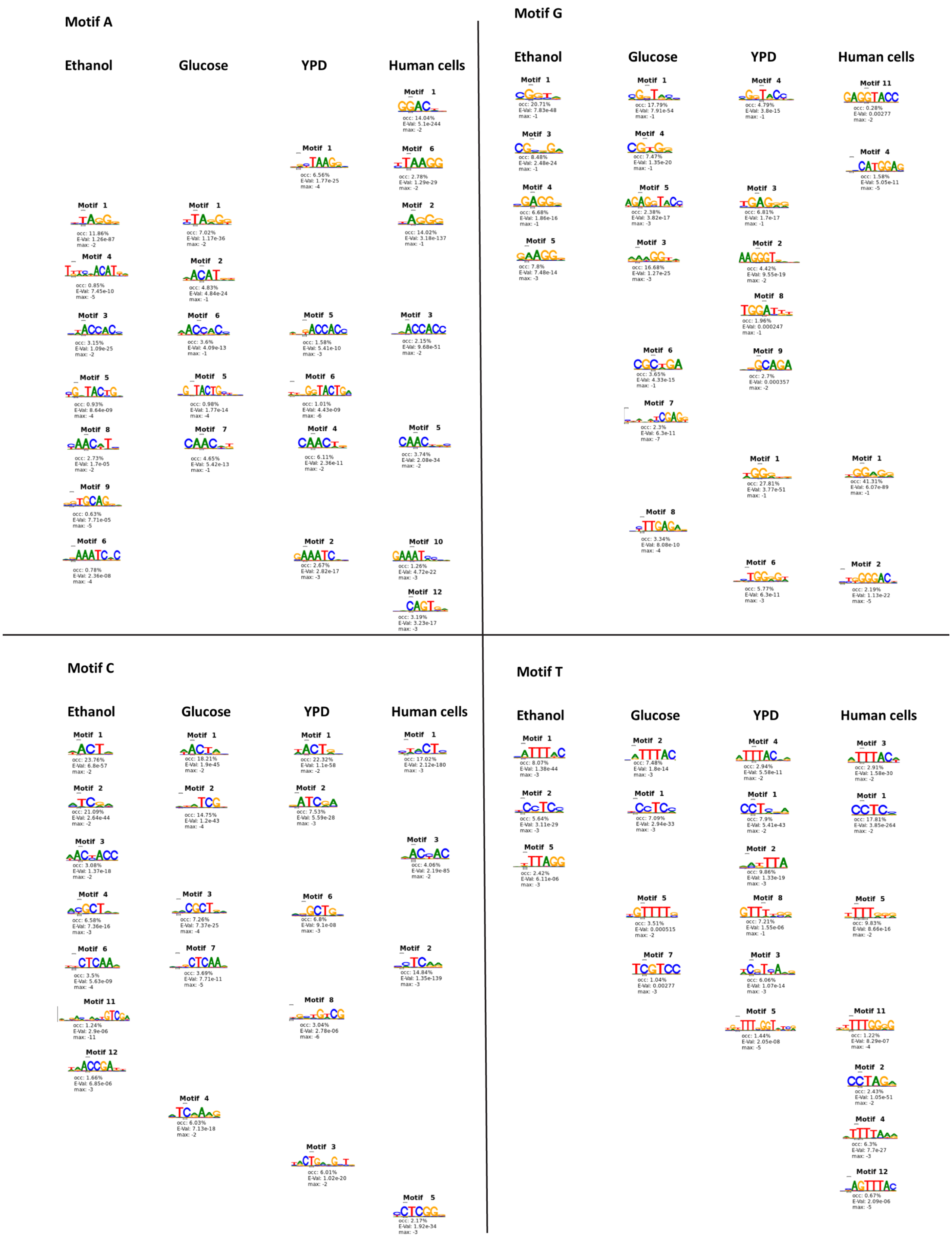


**Supplementary Figure S5.** The high confident identified motifs derived from differential %ESB loci of individual base using XXmotif software as describe by material and method. Uracil is represented by 'T'.

(A)

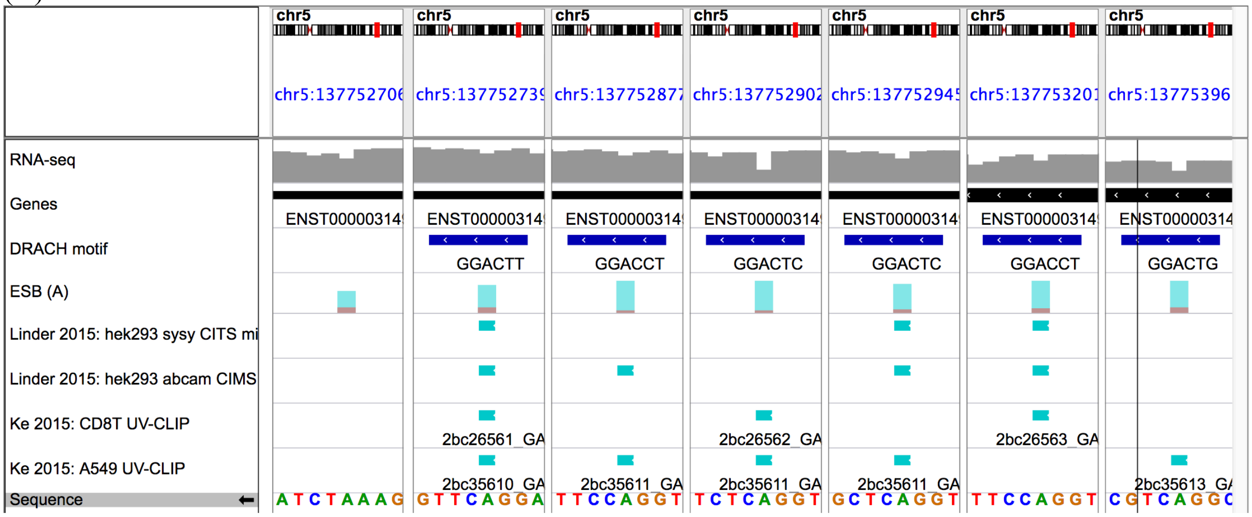


(B)

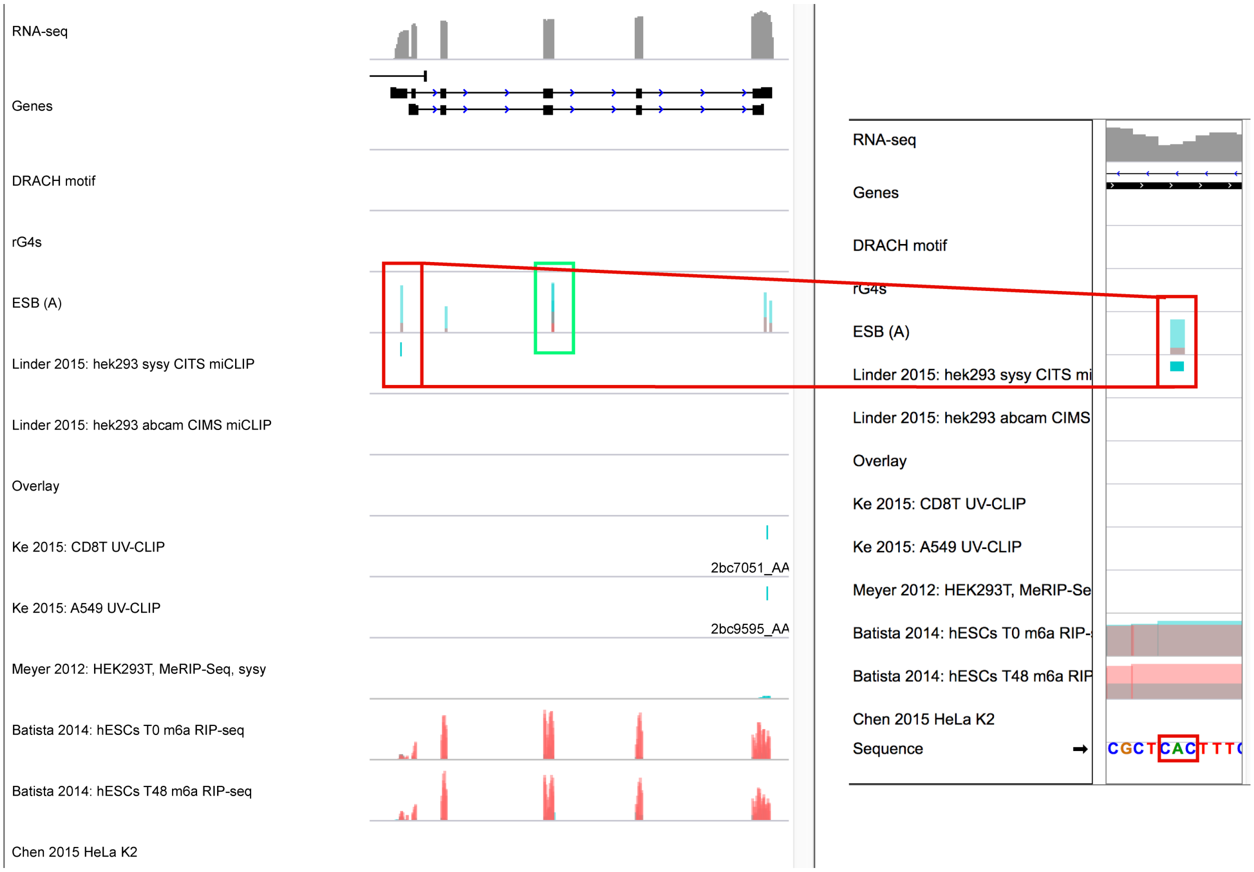


**Supplementary Figure S6** (A) Zoom views of Figure 6D showing the m6A methylations with DRACH motifs that were captured by ELIGOS are consistent at the base resolution level with miCLIPs and UV-CLIPs experimental data as described by Linder *et al*. (*22*) and Ke et al. (*27*). (B) An example of an m6A methylation position detected by the miCLIP method and ELIGOS as shown in the red box. That position contained the artifactual triplet CAC so that it was filtered out from the ELIGOS results (only one position in the green box contains no artifactual triplets that would not be filtered out from the final ELIGOS results).

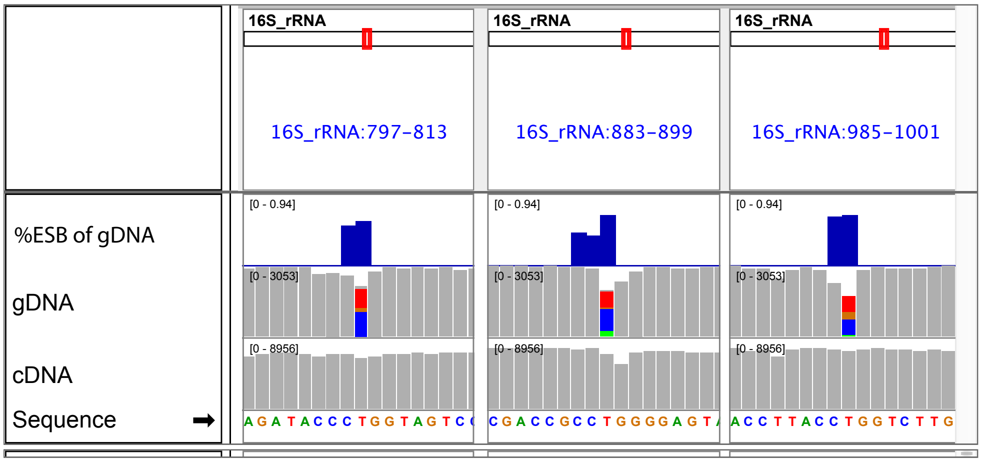


**Supplementary Figure S7**. Examples showing ELIGOS can be applied to identify methylations of genomic DNA (native RNA) coding for rRNA of *E. coli*, shown in IGV genome browser. The native gDNA was compared with cDNA obtained from dcDNA-seq. The motif CCTGG known to be methylated by Dcm methyltransferase [PMC4231299] was correctly identified by ELIGOS.

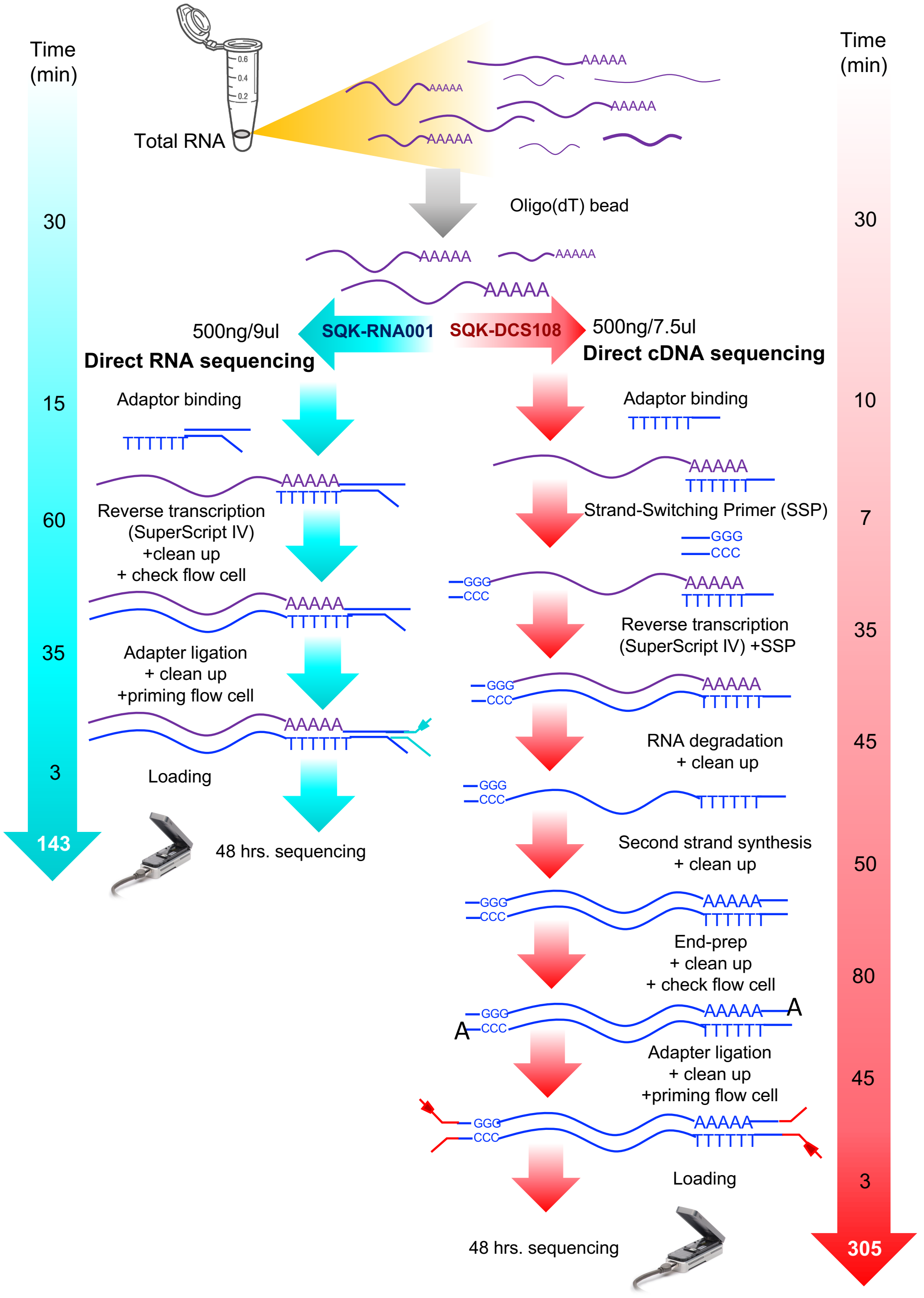


**Supplementary Figure S8.** Summary of the experimental steps involved in library preparation for dRNA-seq (dRNA, left) and dcDNA-seq (dcDNA, right).
